# The novel ECM protein SNED1 mediates cell adhesion via the RGD-binding integrins α5β1 and αvβ3

**DOI:** 10.1101/2024.08.07.606706

**Authors:** Dharma Pally, Nandini Kapoor, Alexandra Naba

## Abstract

The extracellular matrix (ECM) is a complex meshwork comprising over 100 proteins. It serves as an adhesive substrate for cells and, hence, plays critical roles in health and disease. We have recently identified a novel ECM protein, SNED1, and have found that it is required for neural crest cell migration and craniofacial morphogenesis during development and in breast cancer, where it is necessary for the metastatic dissemination of tumor cells. Interestingly, both processes involve the dynamic remodeling of cell-ECM adhesions via cell surface receptors. Sequence analysis revealed that SNED1 contains two amino acid motifs, RGD and LDV, known to bind integrins, the largest class of ECM receptors. We thus sought to investigate the role of SNED1 in cell adhesion. Here, we report that SNED1 mediates breast cancer and neural crest cell adhesion via its RGD motif. We further demonstrate that cell adhesion to SNED1 is mediated by the RGD integrins α5β1 and αvβ3. These findings are a first step toward identifying the signaling pathways activated downstream of the SNED1-integrin interactions guiding craniofacial morphogenesis and breast cancer metastasis.

**SUMMARY STATEMENT:** We report that the novel extracellular matrix protein SNED1 promotes the adhesion of breast cancer cells and neural crest cells via interaction with α5β1 and αvβ3 integrins, the first SNED1 receptors identified to date.

## INTRODUCTION

The extracellular matrix (ECM), a fundamental component of multicellular organisms, is a complex 3-dimensional (3D) meshwork consisting of over a hundred proteins (Hynes and Naba, 2012; Naba, 2024). The primary function of the ECM is to serve as a substrate for cell adhesion (Hynes, 2009). The adhesion of cells to their surrounding ECM is mediated by cell-surface receptors and is critical for cell survival, as detachment from the ECM results in apoptotic cell death, known as anoikis (Frisch and Francis, 1994). In addition, cell-ECM interactions trigger molecular events that regulate a multitude of cellular phenotypes including migration (Dzamba and DeSimone, 2018; Pally and Naba, 2024), proliferation (Hynes, 2009), and differentiation (Walma and Yamada, 2020). As a result, alterations of cell-ECM adhesions and downstream signaling pathways lead to developmental defects (Rozario and DeSimone, 2010) and pathologies like cancer (Cox, 2021; Pickup et al., 2014) and fibrosis (Herrera et al., 2018). Yet, only a small subset of the hundreds of proteins comprising the matrisome is the focus of active investigations, and the mechanisms by which they interact with cells and guide cell phenotype are known for an even smaller subset.

One such understudied ECM protein is Sushi, Nidogen, and EGF like Domains 1 (SNED1). The murine gene *Sned1* was cloned two decades ago (Leimeister et al., 2004), however, it took ten years to identify its first function as a promoter of breast cancer metastasis (Naba et al., 2014). Beyond its role in breast cancer, we recently reported that *Sned1* is an essential gene, as knocking it out resulted in early neonatal lethality and severe craniofacial malformations (Barqué et al., 2021). We further showed that knocking out *Sned1* specifically from neural crest cells, the cell population that contributes to forming most craniofacial features (Mankarious and Goudy, 2010; Martik and Bronner, 2021; Trainor, 2005), was sufficient to recapitulate the craniofacial phenotype observed upon global *Sned1* deletion, demonstrating a new role for SNED1 in craniofacial morphogenesis (Barqué et al., 2021). However, as of today, the mechanisms through which SNED1 interacts with cells to mediate its phenotypes remain unknown. Of note, metastatic breast cancer cells and neural crest cells share common features (Gallik et al., 2017), including their ability to remodel their adhesions to acquire increased migratory potential (Doyle et al., 2022; Gallik et al., 2017; Mayor and Theveneau, 2013; Tucker et al., 1988). This process is, in part, mediated by integrins, which are the main class of ECM receptors (Bökel and Brown, 2002; Campbell and Humphries, 2011; Hood and Cheresh, 2002; Hynes, 2002; Kanchanawong and Calderwood, 2023). For example, *in vivo* and *in vitro* experiments have demonstrated that β1 integrins at the surface of neural crest cells interact with the ECM and ECM proteins like fibronectin to promote cell adhesion and subsequent migration (Alfandari et al., 2003; Duband et al., 1991; Leonard and Taneyhill, 2020; Pietri et al., 2004; Testaz et al., 1999; Yang et al., 1993). β1-containing integrin heterodimers expressed by breast cancer cells have been shown to interact with the vascular ECM to promote extravasation during metastasis (Chen et al., 2016) and be essential to every step of the metastatic cascade (Hamidi and Ivaska, 2018). Interestingly, SNED1 contains two putative integrin-binding motifs, an arginine-glycine-aspartic acid (RGD) triplet and a leucine-aspartic acid-valine (LDV) triplet, known in other proteins to mediate cell-ECM adhesion. We thus sought to determine whether SNED1 played a role in cell adhesion.

Here we report that SNED1 mediates the adhesion of breast cancer cells and neural crest cells, two cell types of relevance to the *in-vivo* functions of SNED1. Using a combination of genetic and pharmacological approaches, we further show that cell adhesion to SNED1 is mediated by its RGD motif and the engagement of α5β1 and αvβ3 integrins. Our study is thus the first to report the identification of SNED1 receptors and constitutes an important step toward the identification of the biochemical signaling events leading to SNED1-dependent breast cancer metastasis and craniofacial development.

## RESULTS AND DISCUSSION

### SNED1 mediates breast cancer and neural crest cell adhesion

We first sought to determine whether SNED1 could mediate cell adhesion. To do so, we seeded highly metastatic MDA-MB-231 ‘LM2’ breast cancer cells (further termed LM2) or O9-1 neural crest cells on surfaces coated with increasing concentrations of purified human SNED1 (Vallet et al., 2021) (Fig S1A) or murine Sned1, respectively, and allowed cells to adhere for 30 min. Cell adhesion was assayed using a crystal-violet-based colorimetric assay and we found that SNED1 mediated the adhesion of LM2 breast cancer cells and O9-1 neural crest cells in a concentration-dependent manner with maximum adhesion observed at 10 μg/mL concentration (Fig 1A and B, respectively). At this concentration, SNED1 exerted the same adhesive properties as fibronectin toward these two cell populations (represented by 100% adhesion based on the standard curve). We further showed that the O9-1 cells, which are of murine origin, adhered in the same proportion to both murine Sned1 and human SNED1 (Fig S1B), perhaps not surprisingly, as the two orthologs share an 84% sequence identity (Barqué et al., 2021).

**Figure 1.**
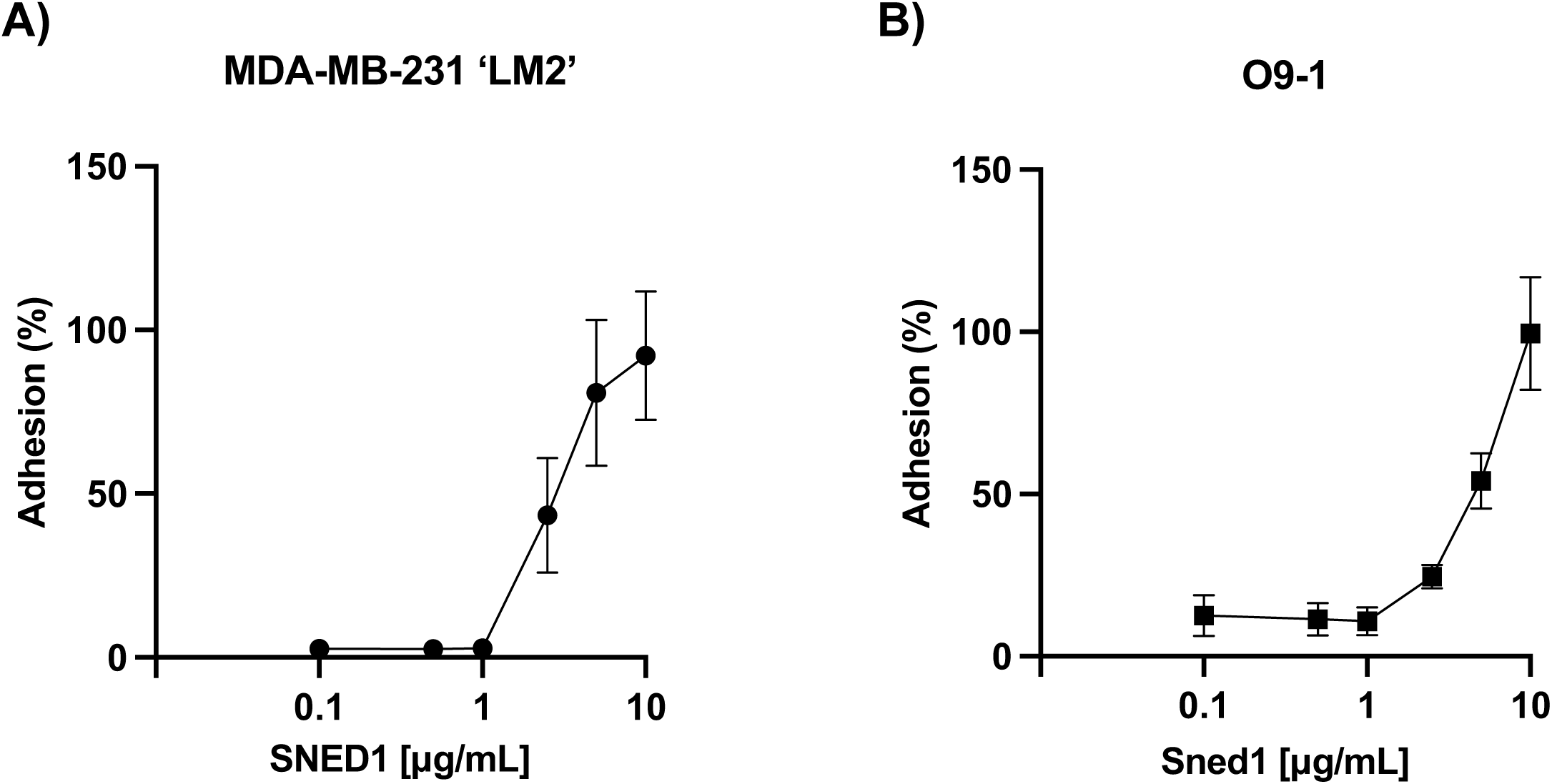
SNED1 promotes breast cancer and neural crest cell adhesion. (A) Adhesion of MDA-MB-231 ‘LM2’ breast cancer cells to purified human SNED1 is concentration-dependent. (B) Adhesion of O9-1 mouse neural crest cells to purified murine Sned1 is concentration dependent. Data is represented as mean ± SD from three biological experiments.

Sned1 also mediated the adhesion of immortalized mouse embryonic fibroblasts derived from *Sned1* knockout mice (*Sned1*^KO^ iMEFs) to the same extent as the other two cell lines (Fig S1C). Altogether these experiments demonstrate that SNED1 mediates cell adhesion and exerts adhesive properties toward a panel of cell types.

### The N-terminal region of SNED1 is sufficient to mediate cell adhesion

SNED1 is a modular protein composed of different domains, including a NIDO domain, a follistatin domain, and a sushi domain (also known as complement control protein or CCP domain), in addition to multiple repeats of EGF-like (EGF), calcium-binding EGF-like (EGF-Ca), and fibronectin type 3 (FN3) domains, all involved in protein-protein interactions (Fig 2A). To assess which region(s) of SNED1 mediate cell adhesion, we generated and purified three truncated forms of SNED1 (Fig 2A): the SNED1^1-751^ form lacks the C-terminal region that comprises three FN3 domains, shown to mediate cell adhesion in other ECM proteins, and the most C-terminal EGF-like domains; the SNED1^1-530^ fragment additionally lacks the sushi domain and EGF-like domains after the follistatin domain; the SNED1^1-260^ construct encompasses the very N-terminal region, including a single NIDO domain, which is only present in four other proteins in the human and mouse proteomes (nidogen-1 and nidogen-2, alpha-tectorin, and mucin-4) and its function remains unknown (Barqué et al., 2021). These synthetic constructs were designed to include adequate domain boundaries compatible with proper folding for the truncated proteins to be secreted (Fig 2B).

**Figure 2.**
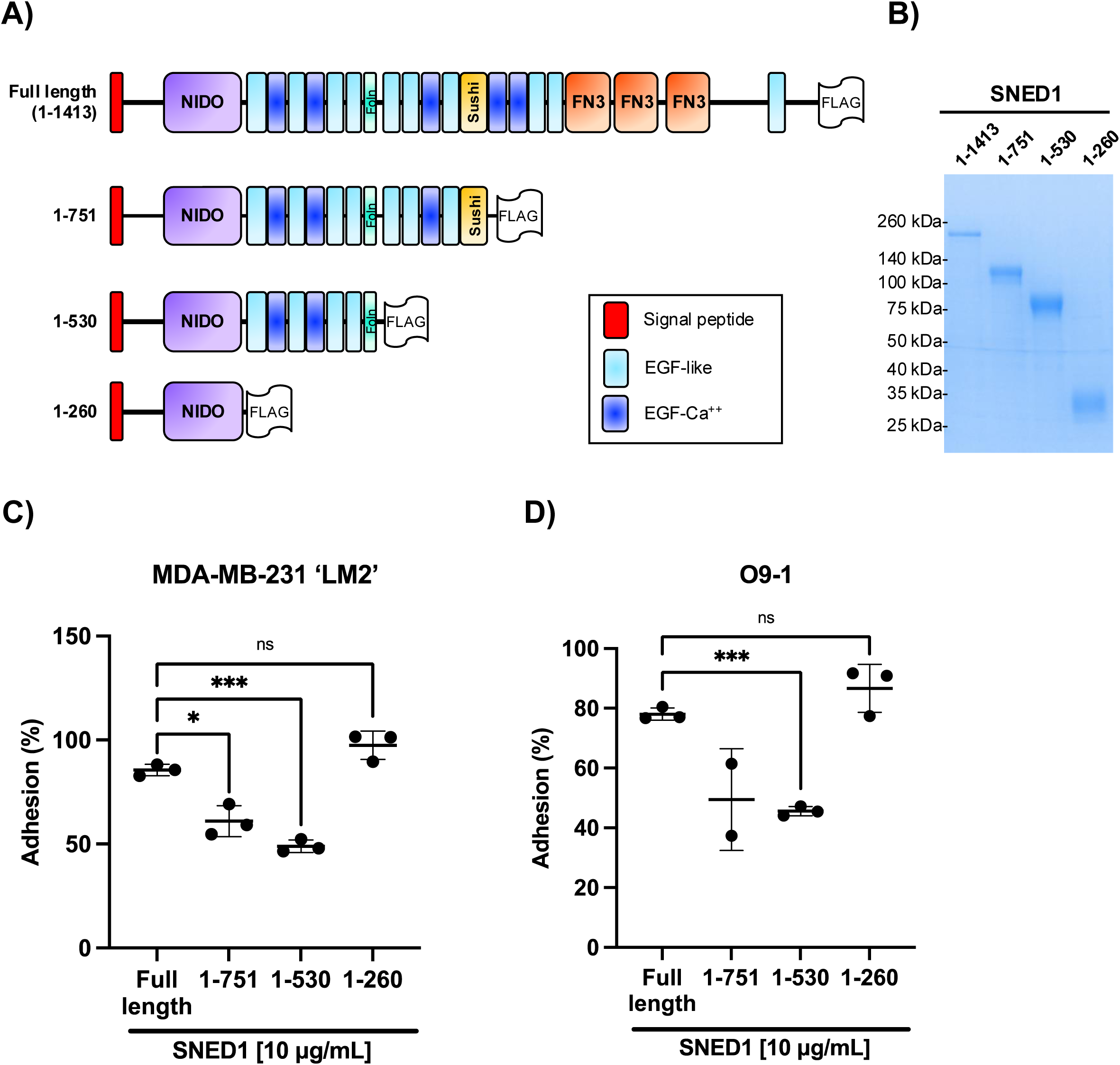
The N-terminal region of SNED1 mediates cell adhesion. (A) Schematic showing FLAG-tagged full-length SNED1 and the different truncated forms of SNED1 used in this study: SNED1^1-751^ encompasses the N-terminal region until the sushi domain; SNED1^1-530^ encompasses the N-terminal region until the follistatin domain; SNED1^1-260^ encompasses the N-terminal region until the NIDO domain. (B) Coomassie-stained gel showing the purity of purified full-length and truncated forms of SNED1. (C-D) Adhesion of MDA-MB-231 ‘LM2’ breast cancer cells (C) and O9-1 neural crest cells (D) to full-length and truncated forms of SNED1. Data is represented as mean ± SD from at least two biological experiments. Welch and Brown-Forsythe one-way ANOVA with Dunnett’s T3 correction for multiple comparisons was performed to determine statistical significance. ns: non-significant, *p<0.05, *** p<0.001.

Using these purified truncated proteins as substrates, we found that LM2 cells had a decreased ability to adhere to SNED1^1-751^ and SNED1^1-530^ as compared to full-length SNED1 (1.4- and 1.75-fold decrease, respectively; Fig 2C). Similarly, we observed that O9-1 cells had a decreased ability to adhere to SNED1^1-751^ and SNED1^1-530^ compared to full-length SNED1 (1.58- and 1.7-fold decrease, respectively; Fig 2D). Notably, although SNED1^1-530^ further lacks the sushi domain and four EGF-like domains, along with the domains absent in SNED1^1-751^, it mediated cell adhesion to the same extent as SNED1^1-751^ for both cell lines (Fig 2C and D). Interestingly, LM2 and O9-1 cells could adhere to the shortest N-terminal SNED1^1-260^ fragment in a similar proportion as to full-length SNED1 (Fig 2C and D, respectively). Since, the SNED1^1-260^ fragment is 6 times smaller than full length SNED1, we thought to perform the same experiment but using similar molar concentration (66.3 μM) rather than amount (10μg/mL), and obtained a similar result, namely that LM2 cell adhesion on SNED1^1-260^ is comparable to full length SNED1 (Fig S2). Altogether, these results indicate that the adhesive property of SNED1 is primarily mediated by its N-terminal region. These results also suggests that the three FN3 domains and the EGF-like domains lacking in the SNED1^1-751^ and SNED1^1-530^ constructs are required for full length SNED1 to adopt a conformation where the N-terminal adhesive site is fully accessible to cells, since their absence resulted in decreased cell adhesion.

### SNED1 mediates cell adhesion via its RGD motif

Analysis of the SNED1 sequence has revealed the presence of two putative integrin binding motifs: RGD and LDV. These motifs were discovered in other ECM proteins, such as fibronectin and thrombospondin 1, and interact with integrin heterodimers at the cell surface to mediate cell adhesion (Lawler et al., 1988; Ruoslahti and Pierschbacher, 1987).

To determine whether the RGD and LDV motifs of SNED1 are required for cell adhesion, we mutated these sites alone or in combination (p40D>E in the RGD motif; p311D>A in the LDV motif; Fig 3A). Similar mutations in other ECM proteins have been shown to disrupt their interaction with integrin heterodimers (Cherny et al., 1993; Pytela et al., 1985). We next expressed these constructs in 293T cells and purified the corresponding secreted proteins from the conditioned culture medium using affinity chromatography (Fig S3). Purified proteins were then used to perform cell adhesion assays. We observed that LM2 cells showed a statistically significant decrease in their ability to adhere to SNED1^RGE^ and SNED1^RGE/LAV^ as compared to SNED1^WT^ (3- and 2.5-fold decrease, respectively; Fig 3B). Similarly, O9-1 cells showed a significant decrease in their ability to adhere to SNED1^RGE^ and SNED1^RGE/LAV^ as compared to SNED1^WT^ (4- and 5.5-fold decrease, respectively; Fig 3C). However, mutation of the LDV motif to LAV did not affect cell adhesion (Fig 3B, C). These results demonstrate that the RGD but not the LDV motif in SNED1 is required for cell adhesion.

**Figure 3.**
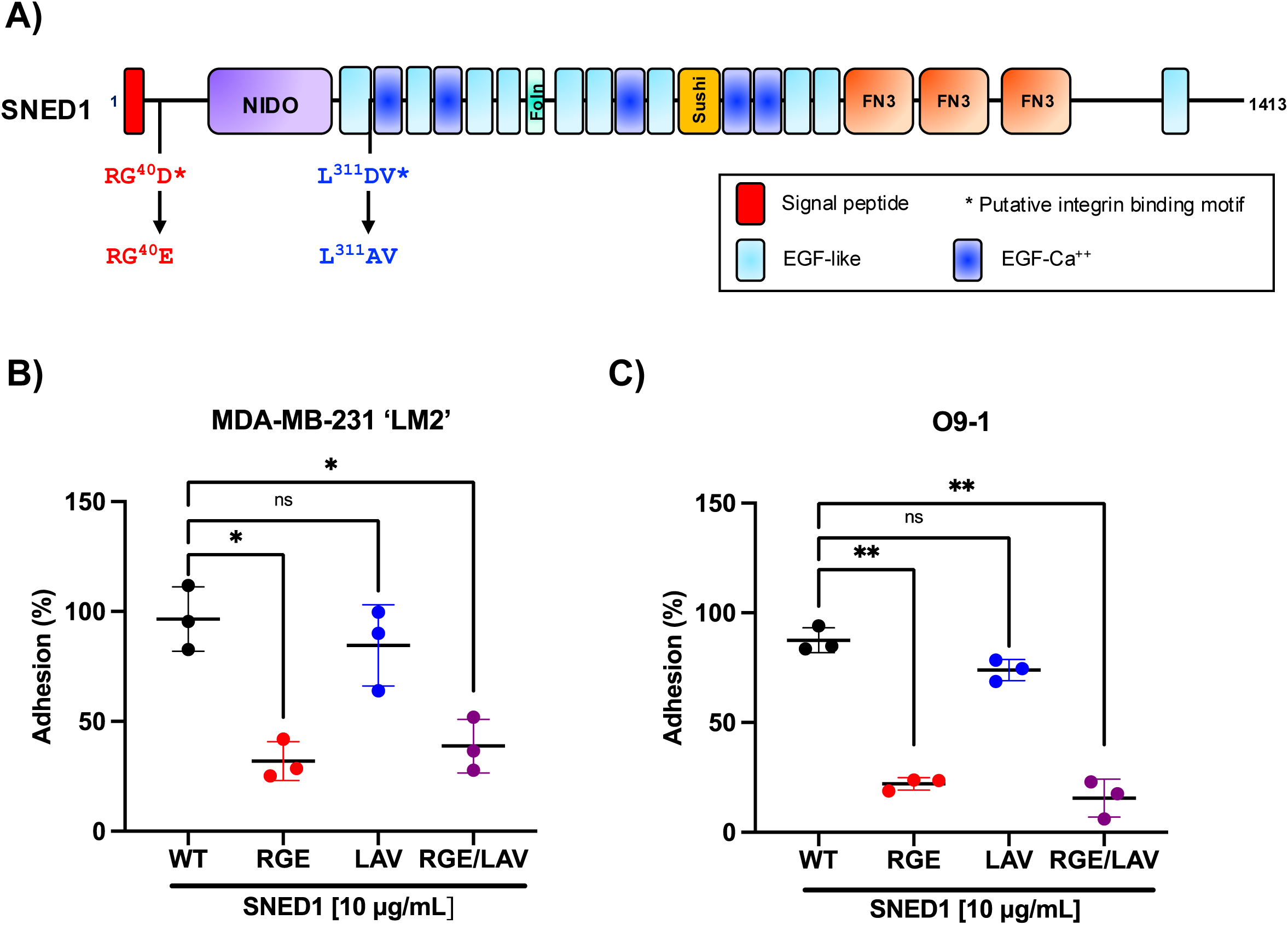
The RGD motif in SNED1 is required to mediate cell adhesion. (A) Schematic of SNED1 showing the two putative integrin-binding motifs in SNED1 (^38^RGD and ^310^LDV) and the point mutations introduced to disrupt their integrin-binding activity (^38^RGD>RGE, ^310^LDV>LAV). (B, C) Adhesion of MDA-MB-231 ‘LM2’ breast cancer cells (B) and O9-1 mouse neural crest cells (C) to wild type or integrin-binding mutants (RGE, LAV or RGE/LAV) of SNED1. Data is represented as mean ± SD from three biological experiments. Welch and Brown-Forsythe one-way ANOVA with Dunnett’s T3 correction for multiple comparisons was performed to determine statistical significance. ns: non-significant, *p<0.05, ** p<0.01.

### Functional inhibition of RGD integrins significantly reduces breast cancer and neural crest cell adhesion to SNED1

To complement this set of observations and determine whether integrins mediate adhesion to SNED1, we performed experiments aimed at targeting the ability of integrins to interact with SNED1. First, we performed adhesion assays in the presence of cyclic RGDfV (cRGDfV) peptide, a peptide known to bind with high affinity to integrins that engage with the RGD motif of ECM proteins (Aumailley et al., 1991). We observed reduced adhesion of both LM2 (Fig 4A) and O9-1 cells (Fig 4B) in a concentration-dependent manner in presence of cRGDfV as compared to vehicle-treated cells. Since we previously showed that the N-terminal region of SNED1 was sufficient to mediate cell adhesion and the RGD motif is located within this fragment, we evaluated the ability of cells to adhere to SNED1^1-260^ in presence of 10μM of cRGDfV peptide. We found that this integrin inhibitor fully abrogated LM2 and O9-1 cell adhesion (Fig S4A, B). This result suggests that the RGD motif at the N-terminal end is essential for the adhesive property of the SNED1^1-260^.

**Figure 4.**
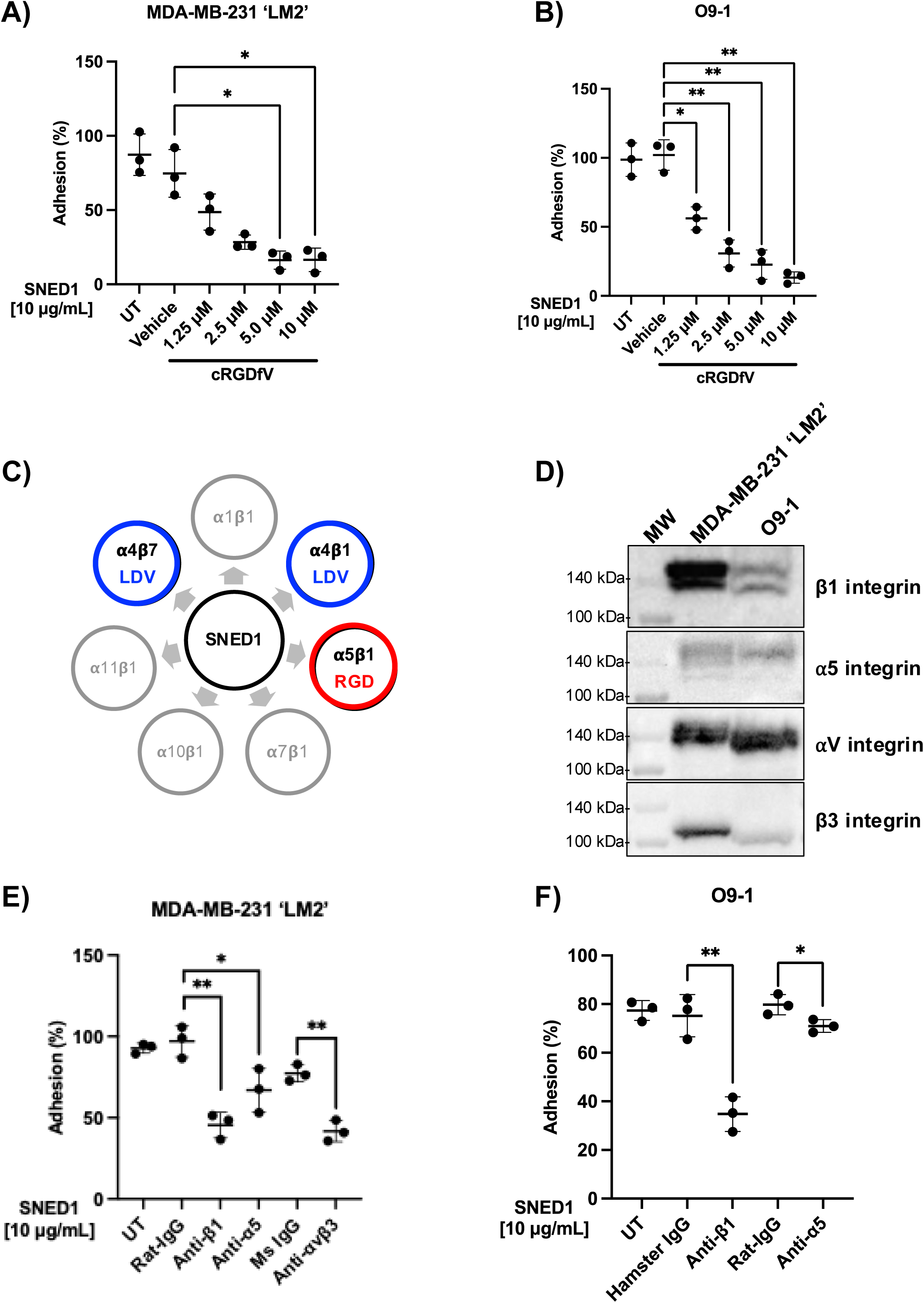
Functional blocking of integrins decreases cell adhesion to SNED1. (A-B) Adhesion of MDA-MB-231 ‘LM2’ breast cancer cell (A) and O9-1 neural crest cells (B) to SNED1 is inhibited in presence of increasing concentrations of cRGDfV peptide. Data is represented as mean ± SD from three biological experiments. Welch and Brown-Forsythe one-way ANOVA with Dunnett’s T3 correction for multiple comparisons was performed to determine statistical significance. *p<0.05, **p<0.01. (C) Schematic showing the seven integrin heterodimers previously predicted to interact with SNED1. Adapted from (Vallet et al., 2021). (D) Immunoblots on total cell extract from MDA-MB-231 ‘LM2’ and O9-1 cells shows β1 integrin, α5 integrin, αv integrin and β3 integrins expression. (E) Adhesion of MDA-MB-231 ‘LM2’ breast cancer cells to SNED1 is decreased in presence of anti-β1, anti-α5 and anti-αVβ3 integrin-blocking antibodies. (F) Adhesion of O9-1 mouse neural crest cells to SNED1 is decreased in presence of anti-β1 and anti-α5 integrin blocking antibodies. Data is represented as mean ± SD from three biological experiments. Unpaired Student’s two-tailed t-test with Welch’s correction was performed to determine statistical significance. *p<0.05, ** p<0.01.

The cRGDfV peptide inhibits a panel of RGD-binding integrin heterodimers such as αvβ3, αvβ6, α5β1, and αvβ5 (Kapp et al., 2017). Using *in silico* molecular modeling, we previously predicted that SNED1 could potentially interact with 11 integrin subunits, α1, α4, α7, α10, α11, β1, β2, β3, β4, β5, and β7 (Vallet et al., 2021) forming six functional heterodimers: α1β1, α4β1, α7β1, α10β1, α11β1, and α4β7 (Hynes, 2002) including the RGD-binding integrin α5β1 (Fig 4C). We thus sought to test whether cell adhesion to SNED1 was dependent on α5β1 integrin. We first confirmed that LM2 and O9-1 cells expressed α5β1 integrin (Fig 4D). We then performed cell adhesion assays in presence of functional blocking antibodies targeting β1 and α5 integrins and demonstrated that the adhesion of LM2 cells (Fig 4E) and O9-1 cells (Fig 4F) to SNED1 was significantly reduced in presence of antibodies targeting β1 integrin (52% and 41% decrease, respectively, as compared to isotype controls) or α5 integrin (30% and 9% decrease, respectively, as compared to isotype controls).

While cRGDfV inhibits α5β1 integrin, αvβ3 is more sensitive to inhibition by this peptide (Gurrath et al., 1992; Pfaff et al., 1994). Since LM2 cells also express αvβ3 (Fig 4D), we sought to test whether αvβ3, could act as a receptor for SNED1, although it was not predicted by our modeling approach. Interestingly, we observed that the adhesion of LM2 cell was significantly decreased in presence of an antibody targeting αvβ3 integrin (35.5% decrease as compared to isotype control; Fig. 4E). This result is in line with our observation of decreased cell adhesion to SNED1^RGE^. Altogether, these experiments identified α5β1 and αvβ3 integrin as the first SNED1 receptors.

It is worth noting that inhibition of α5, β1, or αvβ3 integrins did not fully abrogate cell adhesion to SNED1, suggesting that there are likely additional integrin (such as the RGD-integrin αvβ1 or the non-RGD integrin α1β1) and non-integrin SNED1 receptors at the surface of these cells. Indeed, we have previously shown using i*n-silico* prediction, that SNED1 could interact with 55 transmembrane proteins, in addition to integrins, including the basal cell adhesion molecule (BCAM) or dystroglycan 1 (DAG1), two known ECM receptors (Vallet et al., 2021). The identification of these receptors at the surface of breast cancer cells and neural crest cells will be the focus of future studies.

While our results demonstrate that the RGD motif mediates breast cancer and neural crest cell adhesion, they also raise questions on the role of the LDV motif in SNED1. The LDV motif is primarily recognized by α4β1 and α4β7 integrins that are mainly expressed by leukocytes. Integrin α4 has been previously demonstrated to play a vital role in neural crest cell migration (Kil et al., 1998) and cancer cell adhesion to the vascular endothelium (Taichman et al., 1991). Here, we show that α4 is expressed by LM2 and O9-1 cells (Fig S5A); however, functional blocking of integrin α4 did not affect the adhesion of LM2 cells to SNED1 (Fig S5B), in line with our observation that these cells could adhere similarly to SNED1^LAV^ and SNED1^WT^. This is also in line with the observation that the two truncated forms of SNED1, SNED1^1-530^ and SNED1^1-751^, which contain the LDV motif, are not sufficient to mediate cell adhesion. This opens the possibility that SNED1, via its LDV motif, could engage other cell populations in other pathophysiological processes that have yet to be discovered.

In addition to mediating cell adhesion, integrin heterodimers are involved in the early steps of ECM assembly; for example, α5β1 integrin is critical for the initiation of fibronectin fibrillogenesis (Mao and Schwarzbauer, 2005). We have previously shown that SNED1 forms fibers in the ECM (Vallet et al., 2021), yet we do not know the mechanisms required for this process. It will thus be interesting to determine if α5β1 integrin also mediates SNED1 fiber assembly.

Finally, given the critical role integrin-ECM interactions play in driving pathological processes, devising therapeutic strategies to prevent or disrupt these interactions is an active area of investigation (Hamidi et al., 2016; Pang et al., 2023; Raab-Westphal et al., 2017). To date, seven small molecule inhibitors and biologics have been successfully marketed, including monoclonal antibodies specifically blocking α4β7 and α4β1 to treat inflammatory disorders such as ulcerative colitis, Crohn’s disease, and multiple sclerosis (Pang et al., 2023; Slack et al., 2022). Our study has thus the potential to pave the way to devise novel therapeutic strategies aimed at targeting SNED1-integrin interaction to prevent breast cancer metastasis.

## MATERIALS AND METHODS

### Plasmid constructs

The cDNA encoding full-length human SNED1 cloned into pCMV-XL5 was obtained from Origene (clone SC315884). 6X-His-tagged SNED1 previously described (Vallet et al., 2021) was subcloned into the bicistronic retroviral vector pMSCV-IRES-Hygromycin between the BglII and HpaI sites. FLAG-tagged SNED1 was previously described (Vallet et al., 2021).

Site-directed mutagenesis to generate SNED1^RGE^ (c.C120A; p.D40E) and SNED1^LAV^ (c.A932C; p.D311A) was performed using the QuikChange kit (Agilent #200519) following the manufacturer’s instructions using the cDNA encoding SNED1 from Origene as a template. To obtain the double mutant SNED1^RGE/LAV^ (c.C120A; p.D40E / c.A932C; p.D311A), we introduced the c.C120A mutation in the c.A932C mutant. These constructs were then subcloned into the bicistronic retroviral vector pMSCV-IRES-Hygromycin between the BglII and HpaI sites, and a 6X-Hix-tag was introduced by PCR in 3’ (C-terminus of the protein).

Truncated forms of human SNED1 were subcloned by PCR to generate the following fragments: SNED1^1-530^, encompassing the N-terminal region of SNED1 until the follistatin domain, and SNED1^1-751^, encompassing the N-terminal region of SNED1 until the sushi domain (Fig 2A). These constructs were then subcloned into the bicistronic retroviral vector pMSCV-IRES-Hygromycin between the BglII and HpaI sites, and a FLAG tag (DYKDDDDK) was added at the C-terminus via PCR as previously described (Vallet et al., 2021). FLAG-tagged SNED1^1-260^, encompassing the very N-terminal region of SNED1 was previously described (Vallet et al., 2021). The sequences of the primers used to introduce point mutations or generate truncated fragments of SNED1 are listed in Supplementary Table 1. All constructs were verified by sequencing.

### Cell culture

#### Cell maintenance

Highly metastatic breast cancer cells, MDA-MB-231 ‘LM2’ (termed LM2 in the manuscript), were kindly gifted by Dr. Joan Massagué (Memorial Sloan Kettering Cancer Center, New York, NY). LM2 cells, human embryonic kidney 293T cells (termed 293T in the manuscript) stably overexpressing different constructs of SNED1, and immortalized mouse embryonic fibroblasts isolated from *Sned1*^KO^ mice (*Sned1*^KO^ iMEFs) (Vallet et al., 2021) were cultured in Dulbecco’s Modified Eagle’s medium (DMEM; Corning, #10-017-CV) supplemented with 10% fetal bovine serum (FBS; Sigma, #F0926) and 2 mM glutamine (Corning, #25-005-CI); this formulation is termed “complete medium” in this manuscript. The O9-1 mouse cranial neural crest cell line (Millipore Sigma, #SCC049) was cultured per the manufacturer’s instructions. Briefly, cells were cultured on dishes coated with Matrigel® (Corning, #356234) prepared at 0.18 mg/mL in 1X Dulbecco’s Phosphate Buffered Saline (D-PBS) containing calcium and magnesium (Cytiva, #SH30264.FS) in presence of complete ES cell medium containing 15% FBS and leukemia inhibitory factor (Millipore Sigma, #ES-101-B) and supplemented with 25 ng/mL fibroblast growth factor-2 (FGF-2, R&D systems, #233-FB). All cell lines were maintained at 37°C in a 5% CO_2_ humidified incubator.

#### Retrovirus production

293T cells were plated at ∼ 30% confluency in a 6-well plate. Cells were transfected the following day using a Lipofectamine 3000 (Invitrogen, #L3000-008) mixture containing 1 μg of a retroviral vector with the construct of interest and 0.5 μg each of a packaging vector (pCL-Gag/Pol) and a vector encoding the VSVG coat protein prepared in Opti-MEM™ (Gibco, #31985070). After 24h, the transfection mixture was replaced with fresh complete medium, and cells were cultured for an additional 24h, after which the conditioned medium containing viral particles was collected, passed through 0.45 μ filter, and then either immediately used for cell transduction or stored at -80°C for later use.

#### Generation of 293T cells stably expressing SNED1 constructs

293T cells were seeded at ∼30% confluency. The following day, cells were transduced with undiluted viral particles-containing conditioned medium. 24h after transduction, the medium was replaced with fresh complete medium, and cells were allowed to grow for another 24h before selection with hygromycin (100 μg/mL). Once stable cell lines were established, we assessed the production and secretion of recombinant proteins using immunoblotting of total cell extract (TCE) and conditioned medium, respectively, using either an anti-SNED1, anti-His or anti-FLAG antibody.

### Protein purification

#### Condition medium collection

293T cells stably expressing constructs of interest were seeded in a 15 cm dish in complete medium and allowed to grow until reaching 100% confluency. The culture medium was aspirated, the monolayer was rinsed with 1X D-PBS containing calcium and magnesium, and the medium was replaced with serum-free DMEM medium supplemented with 2 mM glutamine for 48h. The serum-free conditioned medium (CM) containing secreted SNED1 was harvested, and cells were allowed to recover for 48h in complete medium before repeating the next cycle. Cells were discarded after five cycles. An EDTA-free protease inhibitor cocktail (0.067X final concentration; Thermo Scientific, #A32955) was added to the CM, and the CM was centrifuged at 4000 rpm for 10 min to remove any cell or cellular debris. Pre-cleared supernatants were collected and stored at -80°C until further processing.

#### Metal affinity purification of His-tagged SNED1 proteins

6X His-tagged SNED1 proteins (wild-type or integrin-binding mutants) were purified via immobilized metal affinity chromatography (IMAC) using an AKTA Pure system for fast protein liquid chromatography (FPLC) at the UIC Biophysics Core Facility. In brief, conditioned medium (CM) containing the secreted protein of interest was thawed at 4°C, concentrated using a 100-kDa protein concentrator with a polyethersulfone membrane (Thermo Scientific, #88537), buffer-exchanged against a binding buffer containing 20 mM Tris, 500 mM NaCl, 20 mM imidazole (pH 7.5), and filtered using a 0.2 μm filter. In parallel, a HisTrap HP column (column volume: 1 mL) was equilibrated with the binding buffer. The filtered CM was applied to the equilibrated HisTrap column to allow protein binding. The column was then washed with 20 column volumes of binding buffer and bound proteins were eluted with a buffer containing 20 mM Tris-HCl, 500 mM NaCl, 5 mM β-mercaptoethanol (pH 7.5), and a stepwise gradient of increasing imidazole concentration (12.5mM, 50mM, 125mM, 250mM, 375mM, and 500mM). The elution fractions were concentrated, and a buffer exchange was performed with HEPES Buffered Saline (HBS; 10 mM HEPES, 150 mM NaCl, pH 7.5).

#### Purification of FLAG-tagged SNED1 proteins

Since we could not achieve sufficient purity of His-tagged truncated forms of SNED1, we cloned FLAG-tagged versions of these proteins for purification (Fig 2B) In brief, conditioned medium containing FLAG-tagged full-length or truncated forms of SNED1 was thawed at 4°C overnight and concentrated using protein concentrators with a 10kDa weight cut-off for the SNED1^1-260^ and SNED1^1-530^ fragments (Thermo Scientific, #88535), a 30kDa cut-off for the SNED1^1-751^ fragment (Thermo Scientific, #88536), and a 100kDa cut-off for full-length SNED1 (Thermo Scientific, #88537). An anti-FLAG resin (Sigma, #A220), containing monoclonal M2 anti-FLAG antibodies coupled to agarose beads was washed with 20 column volumes of HBS twice. Concentrated CM was applied to the resin and incubated overnight under constant rotation at 4°C to allow protein binding. The following day, the unbound fraction was collected, and the resin was washed with 20 column volumes of HBS thrice. Bound FLAG-tagged proteins were eluted by competing with 200 μg/mL of FLAG peptide in HBS in a stepwise manner, resulting in 4 elution fractions. Protein fractions were then pooled, and buffer-exchanged with HBS using protein concentrators of appropriate molecular weight cut-off (see above) to remove the excess of unbound FLAG peptide.

#### Protein quantification and quality assessment

Purified proteins were quantified by measuring the absorbance at λ=280 nm using a NanoDrop spectrophotometer. To assess their quality and purity, proteins were resolved by electrophoresis on polyacrylamide gels. Gels were stained overnight using Coomassie-based AquaStain (Bulldog, #AS001000) and imaged with a ChemiDoc MP™ imaging system (Bio-Rad).

### Adhesion assay

#### Substrate coating

Adhesion assays were performed as described previously (Humphries, 1998). In brief, wells of a 96-well plate were coated with 50 μL of different concentrations, ranging from 0.1 μg/mL to 10 μg/mL, of purified human SNED1 proteins (full-length, fragments, or integrin-binding mutants) or murine Sned1 (R&D systems, 9335-SN) for 180 min at 37°C. Fibronectin (5 μg/mL; Millipore, #FC010) was used as a positive control. The adsorbed protein was immobilized with 0.5% (v/v) glutaraldehyde for 15 min at room temperature (RT). To prevent cell adhesion to plastic, wells were blocked using 200 μL of 10 mg/mL heat-denatured bovine albumin serum (BSA; Sigma, #A9576). Uncoated wells blocked with BSA (10 mg/mL) alone were used as a negative control.

#### Cell seeding and crystal violet assay

Single-cell suspension of LM2 or O9-1 cells containing 5 x 10^5^ cells/mL were prepared in complete medium. 25,000 cells were added to each well and allowed to adhere for 30 min at 37°C. Loosely attached or non-adherent cells were removed by gently washing the wells with 100 μL of D-PBS^++^ thrice and fixed using 100 μL of 5% (v/v) glutaraldehyde for 30 minutes at RT. Cells were stained with 100 μL of 0.1% (w/v) crystal violet in 200 mM 2-(N-morpholino)ethanesulfonic acid (MES), pH 6.0 for 60 min at RT. After washing the excess of unbound crystal violet, the dye was solubilized in 10% (v/v) acetic acid. Absorbance values were measured at λ = 570 nm using a Bio-Tek Synergy HT microplate reader. Cell adhesion was determined by interpolating the absorbance values from a standard curve that was generated by seeding cells at several dilutions (10%-100%) from the single cell suspension on poly-L-lysine (0.01% w/v; Sigma, P4707) coated wells and fixed directly by adding 5% (v/v) glutaraldehyde.

Integrin-blocking experiments were performed by seeding cells on a SNED1 substrate (10 μg/mL) in presence of a cyclic RGDfV peptide (cRGDfV; Sigma, #SCP0111) at concentrations ranging between 1.25 μM and 10 μM or in presence of 10 μg/mL of anti-β1, anti-α5, anti-α4, or anti-αvβ3 integrin-blocking antibodies (see Table S2 for detailed description of all antibodies used in this study).

### Immunoblotting

LM2 and O9-1 cells were cultured for three days in complete medium and lysed in 3x Laemmli buffer (0.1875 M Tris-HCl, 6% SDS, 30% glycerol) containing 100 mM dithiothreitol. Cell lysates were passed through a 26^1/2^-gauge needle to ensure complete cell lysis, and samples were heated at 95°C for 10 min. Lysates were resolved by gel electrophoresis on polyacrylamide gels at constant current (20 mA for stacking, 25 mA for resolving). Proteins were transferred onto nitrocellulose membranes at constant voltage (100 V) for 180 min at 4°C. Membranes were incubated in 5% (w/v) non-fat milk prepared in 1X PBS + 0.1% Tween-20 (PBST) for 60 min at room temperature to prevent non-specific antibody binding and then incubated in the presence of primary anti-β1 integrin, anti-α5 integrin, anti-αV integrin, anti-β3 integrin or anti-α4 integrin antibodies (see Table S2 for details) in 5% (w/v) non-fat milk in PBST overnight at 4°C. Membranes were washed and incubated with horseradish peroxidase (HRP)-conjugated secondary antibodies for 60 min at RT. Immunoreactive bands were detected using chemiluminescence (Thermo Scientific, #E32109) and imaged with a ChemiDoc MP™ imaging system (Bio-Rad).

### Data analysis and statistics

All experiments were performed with at least three technical replicates (n) and, unless noted otherwise, repeated independently three times (N; biological replicates). Data is represented as a mean±standard deviation from three biological replicates. Unpaired Student’s two-tailed *t*-test with Welch’s correction or Welch and Brown-Forsythe one-way ANOVA with Dunnett’s T3 correction for multiple comparisons was performed to measure statistical significance. Plots were generated using PRISM (GraphPad).

## ACKNOWLEDGEMENTS

We would like to thank Dr. Hyun Lee, director of the Biophysics Core facility at UIC, and her team for their help with protein purification. We would also like to thank Dr. Sandra Pinho for recommendations on anti-α4 integrin antibodies and all the members of the Naba lab for insightful discussions.

## COMPETING INTERESTS

The Naba laboratory holds a sponsored research agreement with Boehringer-Ingelheim for work not related to the content of this manuscript. AN holds consulting agreements with AbbVie, XM Therapeutics, and RA Capital.

## FUNDING

This work was partly supported by the National Institute of General Medical Sciences of the National Institutes of Health [R01CA232517], an award from the UIC Chancellor’s Translational Research Initiative, and a start-up fund from the Department of Physiology and Biophysics of the University of Illinois Chicago to AN. NK was supported by an award from the UIC Liberal Arts and Sciences Undergraduate Research Initiative (LASURI) and a UIC Honors College Undergraduate Research Grant.

## DATA AVAILABILITY

All relevant data can be found within the article and its supplementary information. Research materials are available upon request to Dr. Naba.

## SUPPLEMENTARY FIGURE LEGENDS

**Figure S1.**
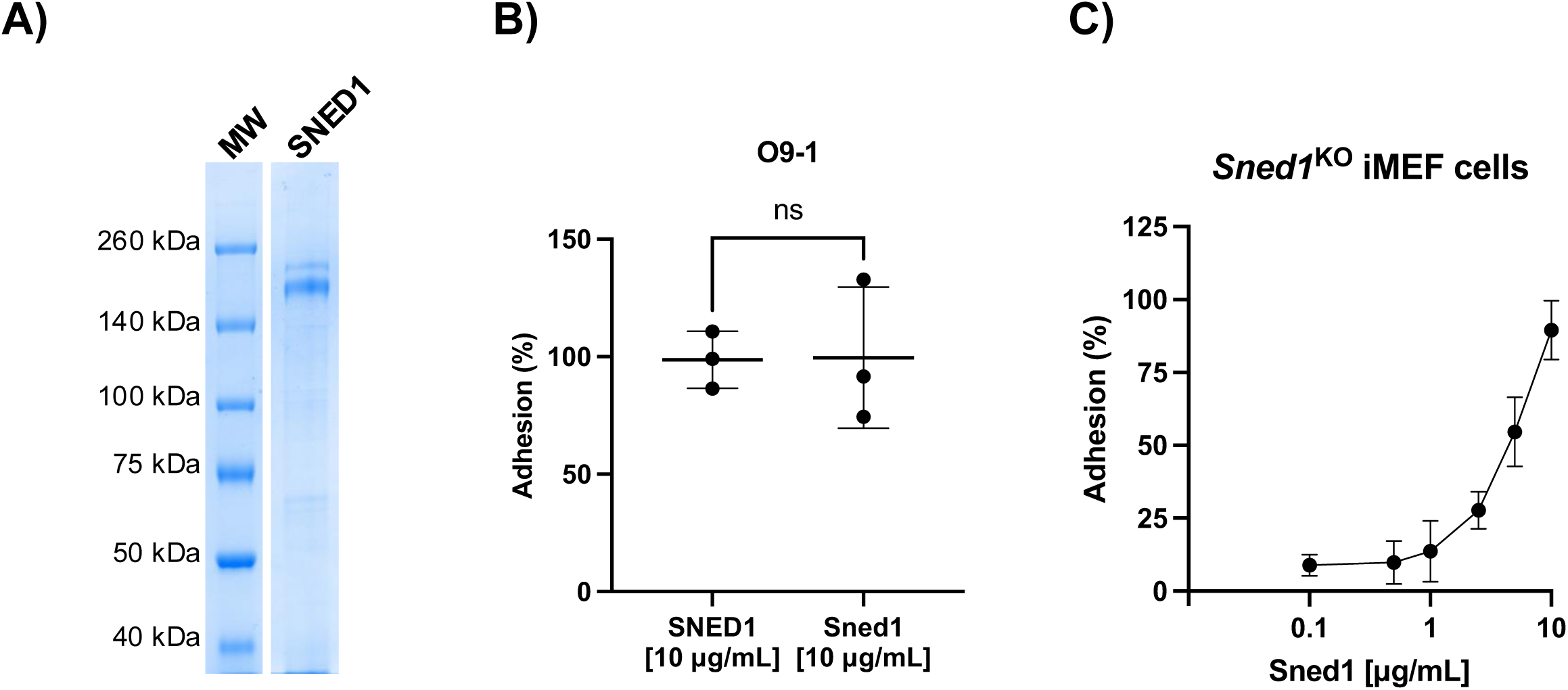
Cell adhesion on human SNED1 and murine Sned1. (A) Coomassie-blue-stained polyacrylamide gel showing the quality and purity of affinity-purified full-length SNED1-His. (B) Murine O9-1 neural crest cells adhere to the same extent to human SNED1 and murine Sned1. Data is represented as mean ± SD from three biological experiments. Unpaired Student’s two-tailed t-test with Welch’s correction was performed to test statistical significance. ns: non-significant. (C) Graph showing the adhesion of immortalized embryonic fibroblast cells isolated from *Sned1* knockout mice (*Sned1*^KO^ iMEF) on increasing concentrations of Sned1.

**Figure S2.**
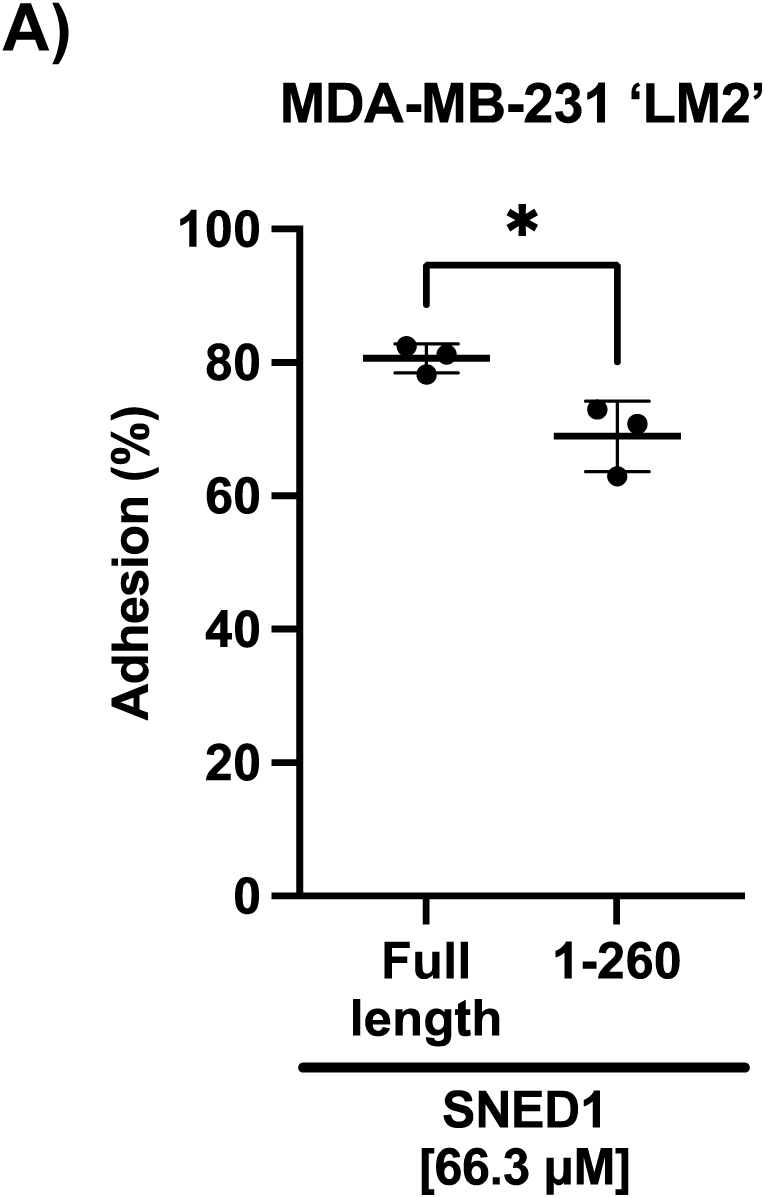
LM2 cell adhesion on equimolar concentration of full length SNED1 and SNED1^1-260^. Graph showing LM2 cell adhesion on 66.3 μM of full length SNED1 and SNED1^1-260^.

**Figure S3.**
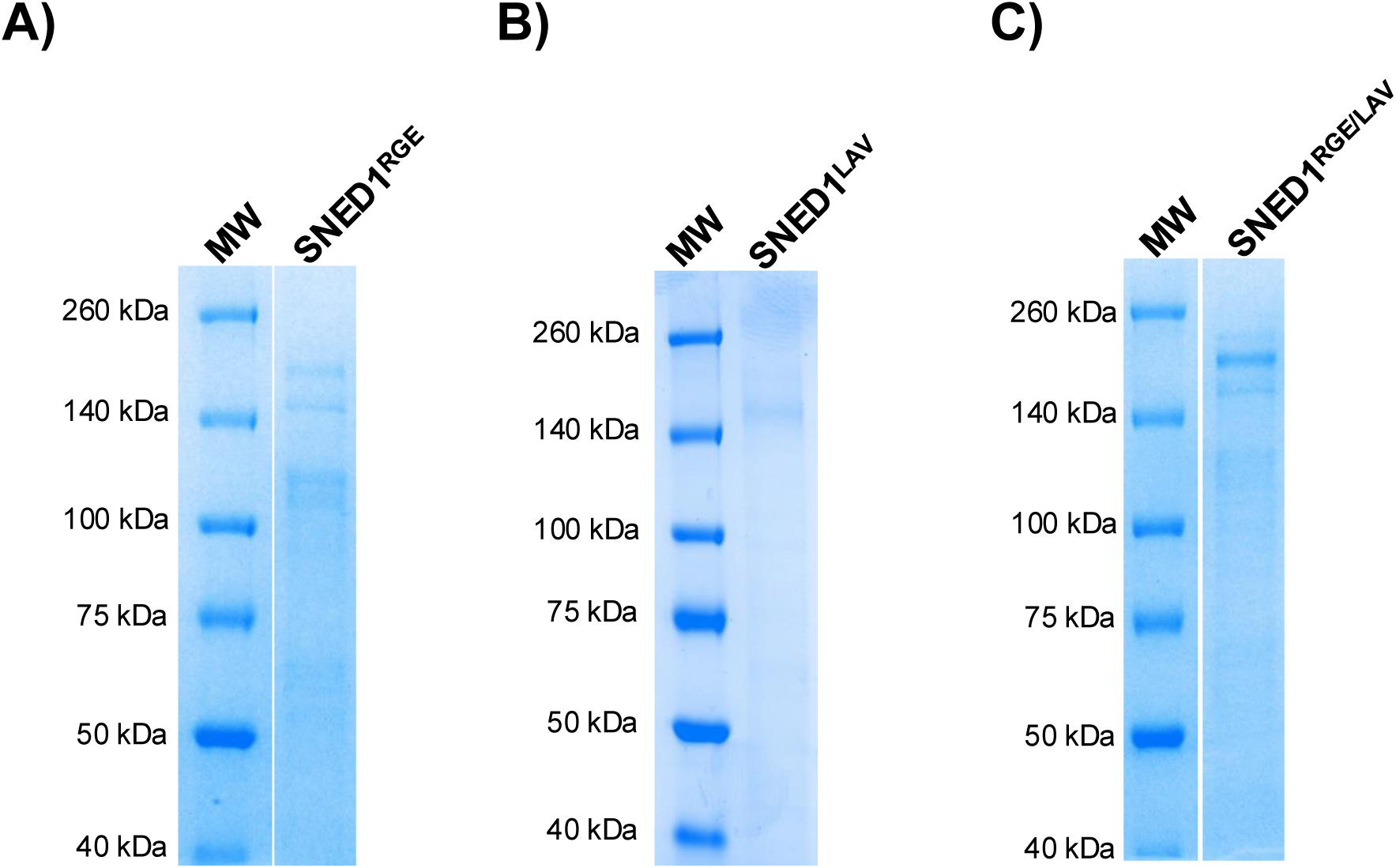
Purification of integrin-binding mutants of SNED1. Coomassie-blue-stained polyacrylamide gels showing the quality and purity of affinity-purified His-tagged SNED1^RGE^ (A), SNED1^LAV^ (B), and SNED1^RGE/LAV^ (C). The different bands correspond to different levels of glycosylation of the proteins, as previously shown (Vallet et al., 2021).

**Figure S4.**
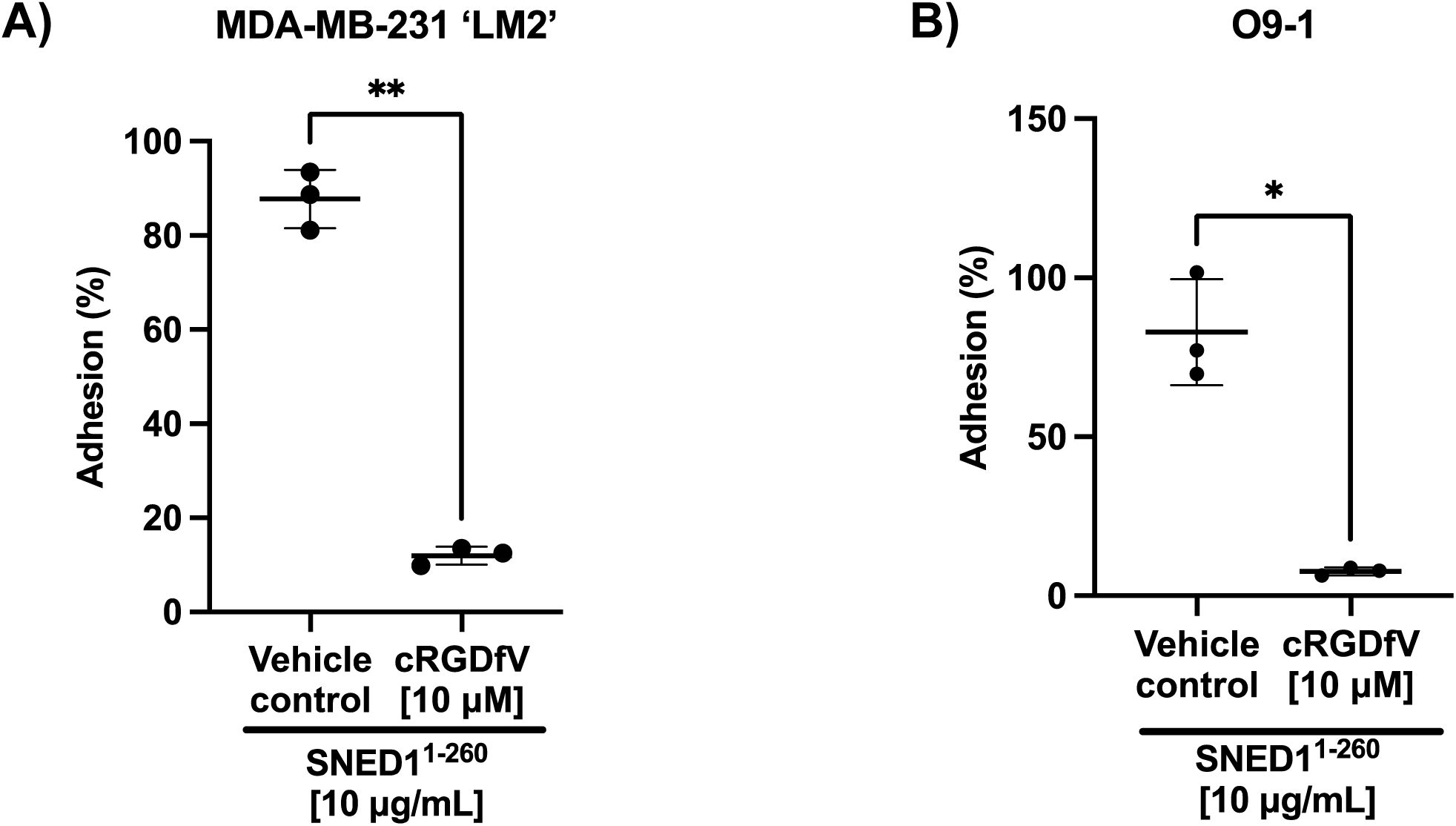
The RGD motif in SNED1^1-260^ is sufficient to mediate cell adhesion. Adhesion of MDA-MB-231’ LM2’ breast cancer cells (A) and O9-1 neural crest cells (B) to the N-terminal fragment of SNED1 (SNED1^1-260^) is significantly decreased in presence of the integrin-binding cRGDfV peptide. Data is represented as mean ± SD from three biological experiments. Unpaired Student’s two-tailed t-test with Welch’s correction was performed to determine statistical significance. *p<0.05, ** p<0.01.

**Figure S5.**
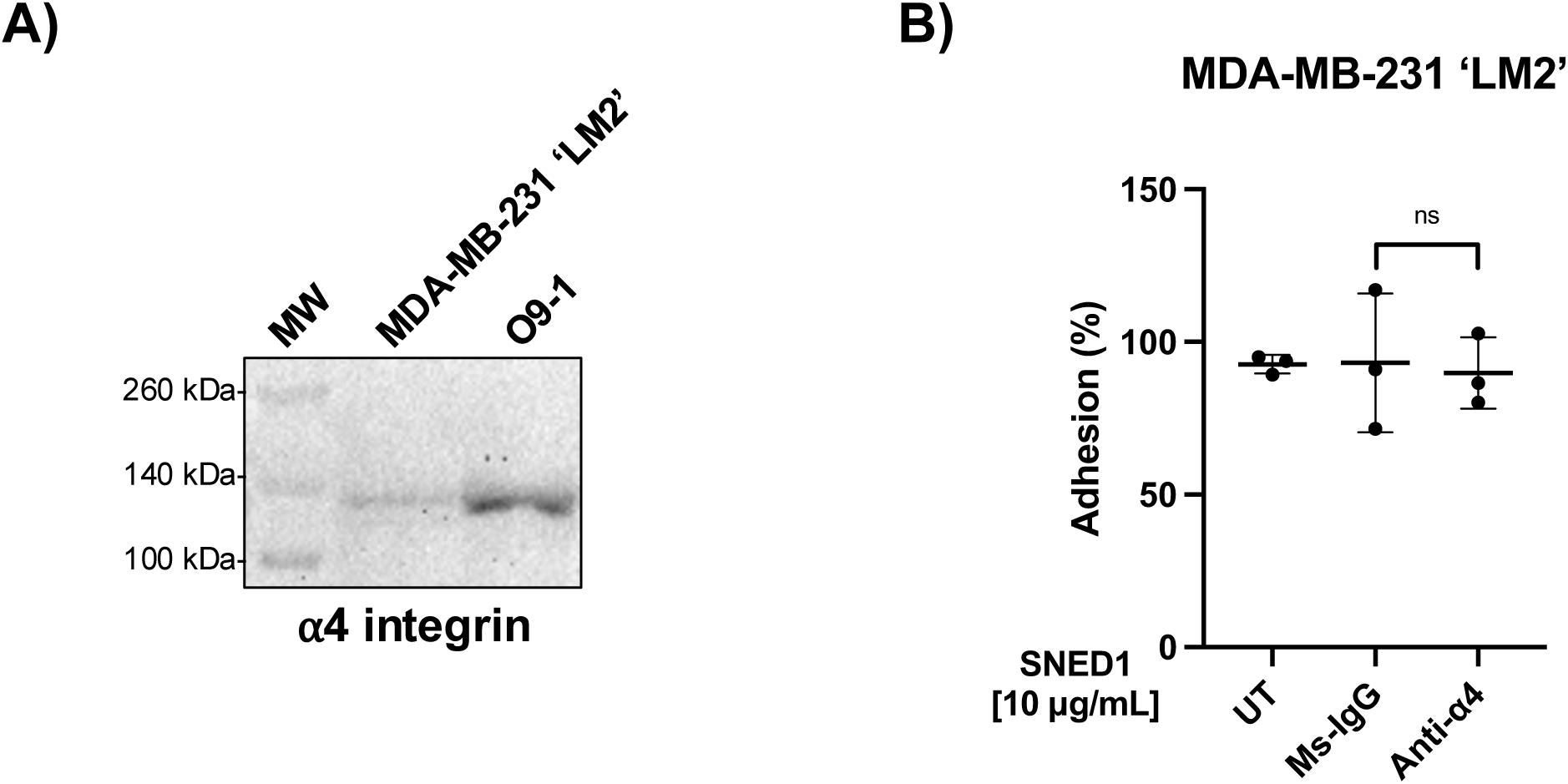
Functional blocking of integrin α4 does not affect breast cancer cell adhesion on SNED1. (A) Immunoblot on total cell extract from MDA-MB-231’ LM2’ and O9-1 cells showing α4 integrin expression. (B) Adhesion of MDA-MB-231’ LM2’ breast cancer cells to SNED1 is not altered in presence of anti-α4 integrin-blocking antibody. Data is represented as mean ± SD from three biological experiments. Unpaired Student’s two-tailed t-test with Welch’s correction was performed to test statistical significance. ns: non-significant.

**Figure S6.**
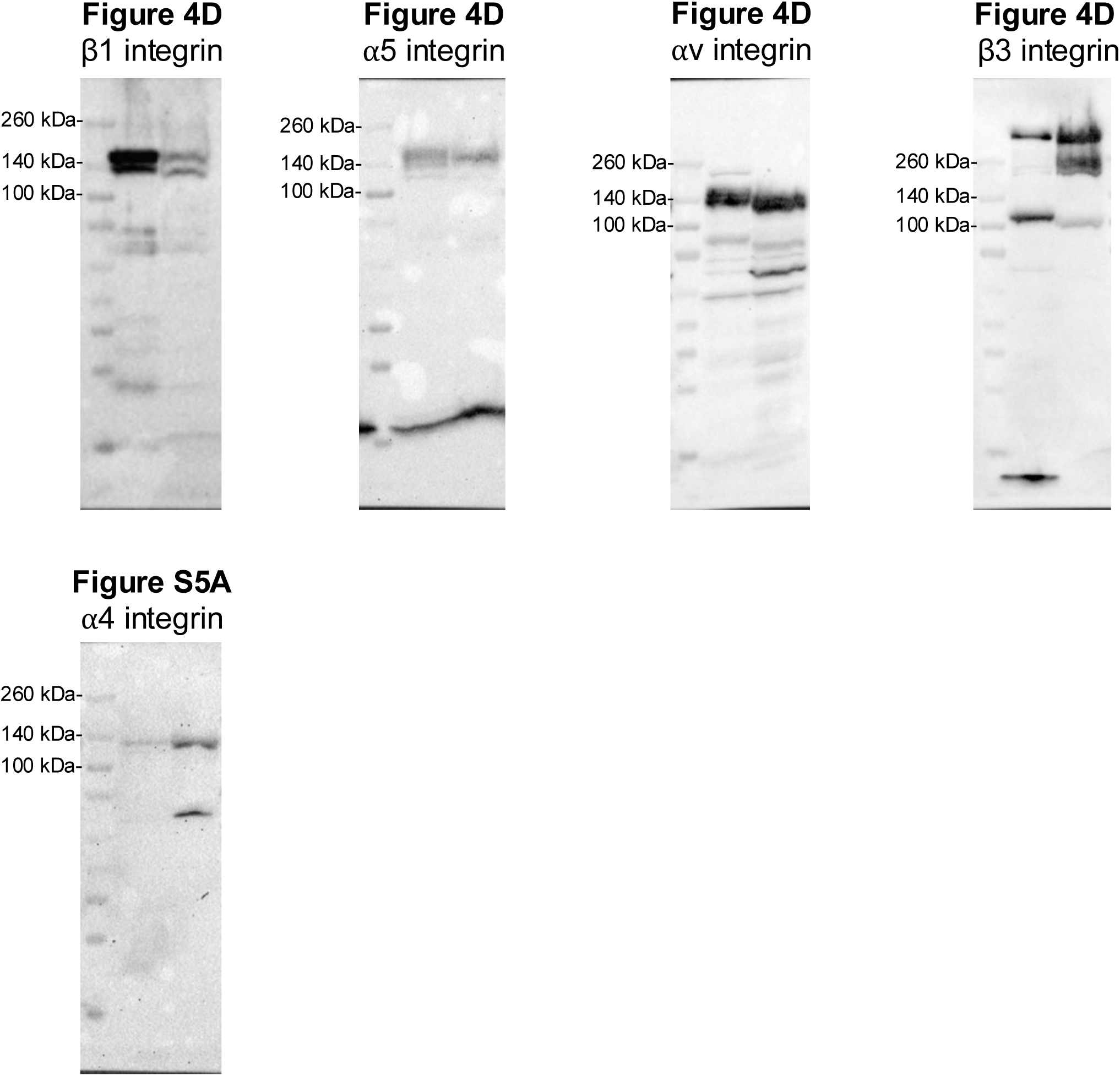
Immunoblot transparency. Uncropped immunoblots for β1 integrin, α5 integrin, αv integrin, β3 integrin, and α4 integrin.

**Supplementary Table 1:**
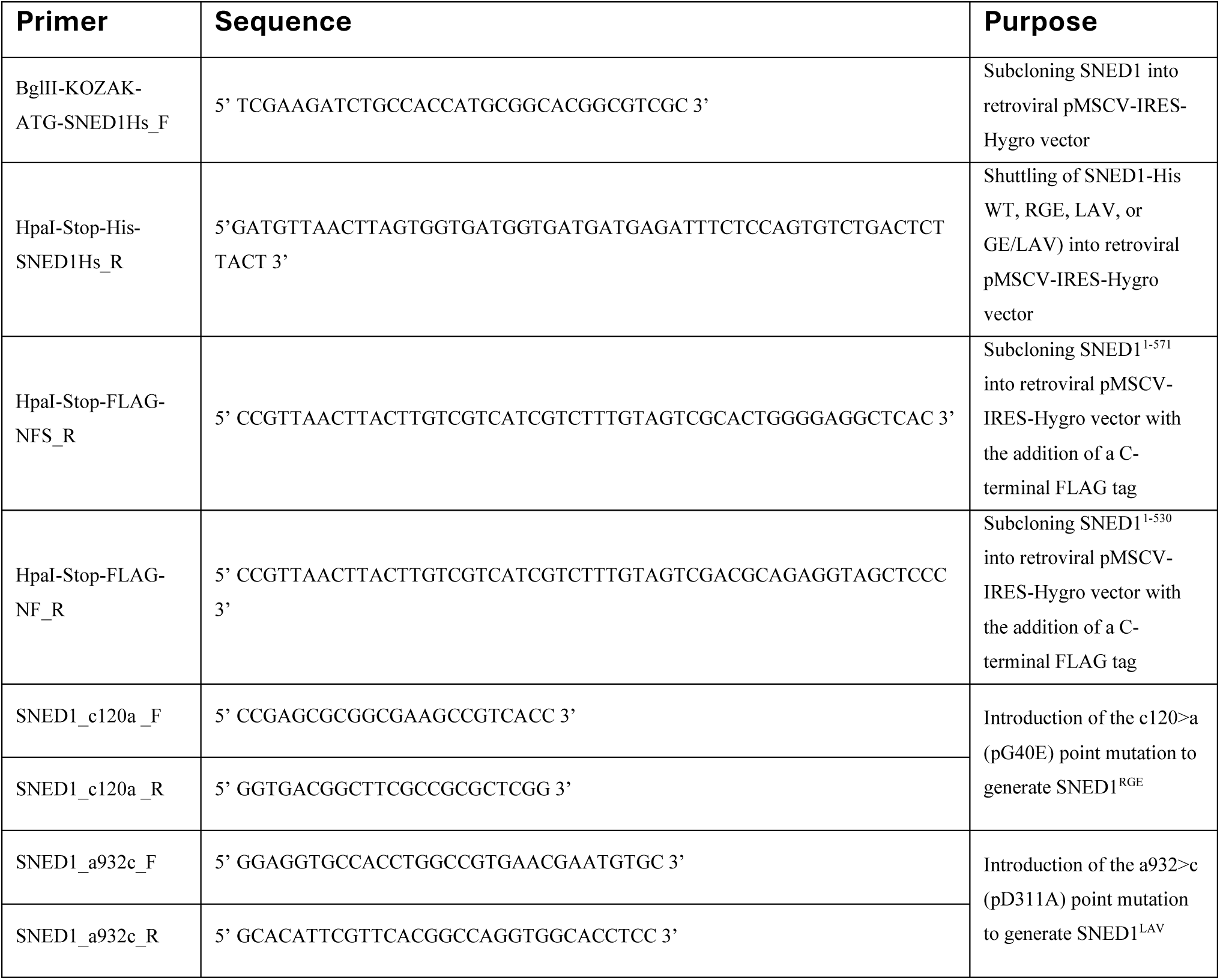
List of primers.

**Supplementary Table 2:** List of antibodies used for functional blocking and immunoblotting.

| Antibody | Host species | Reactivity | Application | Concentration used (µg/mL) | Catalog # |
| --- | --- | --- | --- | --- | --- |
| Anti- β1 Integrin | Rat | Human | Functional blocking | 10 | Sigma, MABT821 |
| Anti-α5 Integrin | Rat | Human | Functional blocking | 10 | Sigma, MABT820 |
| Anti-α4 Integrin | Mouse | Human | Functional blocking | 10 | Sigma, MAB1383 |
| Anti-β1 Integrin | Hamster | Mouse | Functional blocking | 10 | BD Biosciences, BDB555002 |
| Anti-α5 Integrin | Rat | Mouse | Functional blocking | 10 | Biolegend, 103817 |
| Anti-αvβ3 Integrin | Mouse | Human | Functional blocking | 10 | Sigma, MAB1976Z |
| Rat IgG | Rat | Isotype control | Functional blocking | 10 | Invitrogen, PI31903 |
| Mouse IgG | Mouse | Isotype control | Functional blocking | 10 | Invitrogen, PI31933 |
| Hamster IgM | Hamster | Isotype control | Functional blocking | 10 | Biolegend, 401014 |
| Anti-SNED1 | Rabbit | Human | Immunoblotting | 1 | Naba lab |
| Anti-His | Mouse | - | Immunoblotting | 1 | Sigma, SAB2702218 |
| Anti-FLAG | Rabbit | - | Immunoblotting | 1 | Sigma, F7425 |
| Anti-β1 Integrin sera | Rabbit | Human, Mouse | Immunoblotting | 1:1000 dilution | In-house |
| Anti-α5 Integrin | Rabbit | Human, Mouse | Immunoblotting | 0.217 | Abcam, AB150361 |
| Anti-α4 Integrin | Rabbit | Human, Mouse | Immunoblotting | 0.5 | Invitrogen, MA5-27947 |
| Anti-αv Integrin | Rabbit | Human, Mouse | Immunoblotting | 1 | Invitrogen, MA5-32195 |
| Anti-β3 Integrin | Rabbit | Human, Mouse | Immunoblotting | 1 | Invitrogen, MA5-32077 |

## Notes

### Summary of Updates

We are now showing that in addition to α5β1 integrin, another RGD-binding integrin, αvβ3, can also mediate the adhesion of breast cancer cells to SNED1.

## REFERENCES

Alfandari, D., Cousin, H., Gaultier, A., Hoffstrom, B. G. and DeSimone, D. W. (2003). Integrin α5β1 supports the migration of *Xenopus* cranial neural crest on fibronectin. Developmental Biology 260, 449–464.

Aumailley, M., Gurrath, M., Müller, G., Calvete, J., Timpl, R. and Kessler, H. (1991). Arg-Gly-Asp constrained within cyclic pentapoptides Strong and selective inhibitors of cell adhesion to vitronectin and laminin fragment P1. FEBS Letters 291, 50–54.

Barqué, A., Jan, K., De La Fuente, E., Nicholas, C. L., Hynes, R. O. and Naba, A. (2021). Knockout of the gene encoding the extracellular matrix protein SNED1 results in early neonatal lethality and craniofacial malformations. Developmental Dynamics 250, 274– 294.

Bökel, C. and Brown, N. H. (2002). Integrins in Development: Moving on, Responding to, and Sticking to the Extracellular Matrix. Developmental Cell 3, 311–321.

Campbell, I. D. and Humphries, M. J. (2011). Integrin Structure, Activation, and Interactions. Cold Spring Harb Perspect Biol 3, a004994.

Chen, M. B., Lamar, J. M., Li, R., Hynes, R. O. and Kamm, R. D. (2016). Elucidation of the Roles of Tumor Integrin β1 in the Extravasation Stage of the Metastasis Cascade. Cancer Research 76, 2513–2524.

Cherny, R. C., Honan, M. A. and Thiagarajan, P. (1993). Site-directed mutagenesis of the arginine-glycine-aspartic acid in vitronectin abolishes cell adhesion. Journal of Biological Chemistry 268, 9725–9729.

Cox, T. R. (2021). The matrix in cancer. Nat Rev Cancer 21, 217–238.

Doyle, A. D., Nazari, S. S. and Yamada, K. M. (2022). Cell-extracellular matrix dynamics. Phys Biol 19,.

Duband, J.-L., Dufour, S., Yamada, S. S., Yamada, K. M. and Thiery, J. P. (1991). Neural crest cell locomotion induced by antibodies to β1 integrins A tool for studying the roles of substratum molecular avidity and density in migration. Journal of Cell Science 98, 517– 532.

Dzamba, B. J. and DeSimone, D. W. (2018). Chapter Seven - Extracellular Matrix (ECM) and the Sculpting of Embryonic Tissues. In Current Topics in Developmental Biology (ed. Litscher, E. S.) and Wassarman, P. M.), pp. 245–274. Academic Press.

Frisch, S. M. and Francis, H. (1994). Disruption of epithelial cell-matrix interactions induces apoptosis. J Cell Biol 124, 619–626.

Gallik, K. L., Treffy, R. W., Nacke, L. M., Ahsan, K., Rocha, M., Green-Saxena, A. and Saxena, A. (2017). Neural crest and cancer: Divergent travelers on similar paths. Mechanisms of Development 148, 89–99.

Gurrath, M., Müller, G., Kessler, H., Aumailley, M. and Timpl, R. (1992). Conformation/activity studies of rationally designed potent anti-adhesive RGD peptides. European Journal of Biochemistry 210, 911–921.

Hamidi, H. and Ivaska, J. (2018). Every step of the way: integrins in cancer progression and metastasis. Nat Rev Cancer 18, 533–548.

Hamidi, H., Pietilä, M. and Ivaska, J. (2016). The complexity of integrins in cancer and new scopes for therapeutic targeting. Br. J. Cancer 115, 1017–1023.

Herrera, J., Henke, C. A. and Bitterman, P. B. (2018). Extracellular matrix as a driver of progressive fibrosis. J Clin Invest 128, 45–53.

Hood, J. D. and Cheresh, D. A. (2002). Role of integrins in cell invasion and migration. Nature Reviews Cancer 2, 91–100.

Humphries, M. J. (1998). Cell-Substrate Adhesion Assays. Current Protocols in Cell Biology 00, 9.1.1–9.1.11.

Hynes, R. O. (2002). Integrins: Bidirectional, Allosteric Signaling Machines. Cell 110, 673–687.

Hynes, R. O. (2009). The Extracellular Matrix: Not Just Pretty Fibrils. Science 326, 1216–1219.

Hynes, R. O. and Naba, A. (2012). Overview of the Matrisome—An Inventory of Extracellular Matrix Constituents and Functions. Cold Spring Harbor Perspectives in Biology 4, a004903.

Kanchanawong, P. and Calderwood, D. A. (2023). Organization, dynamics and mechanoregulation of integrin-mediated cell–ECM adhesions. Nat Rev Mol Cell Biol 24, 142–161.

Kapp, T. G., Rechenmacher, F., Neubauer, S., Maltsev, O. V., Cavalcanti-Adam, E. A., Zarka, R., Reuning, U., Notni, J., Wester, H.-J., Mas-Moruno, C., et al. (2017). A Comprehensive Evaluation of the Activity and Selectivity Profile of Ligands for RGD-binding Integrins. Sci Rep 7, 39805.

Kil, S. H., Krull, C. E., Cann, G., Clegg, D. and Bronner-Fraser, M. (1998). The α4Subunit of Integrin Is Important for Neural Crest Cell Migration. Developmental Biology 202, 29– 42.

Lawler, J., Weinstein, R. and Hynes, R. O. (1988). Cell attachment to thrombospondin: the role of ARG-GLY-ASP, calcium, and integrin receptors. J Cell Biol 107, 2351–2361.

Leimeister, C., Schumacher, N., Diez, H. and Gessler, M. (2004). Cloning and expression analysis of the mouse stroma marker Snep encoding a novel nidogen domain protein. Developmental Dynamics 230, 371–377.

Leonard, C. E. and Taneyhill, L. A. (2020). The road best traveled: Neural crest migration upon the extracellular matrix. Semin Cell Dev Biol 100, 177–185.

Mankarious, L. A. and Goudy, S. L. (2010). Craniofacial and upper airway development. Paediatric Respiratory Reviews 11, 193–198.

Mao, Y. and Schwarzbauer, J. E. (2005). Fibronectin fibrillogenesis, a cell-mediated matrix assembly process. Matrix Biology 24, 389–399.

Martik, M. L. and Bronner, M. E. (2021). Riding the crest to get a head: neural crest evolution in vertebrates. Nat Rev Neurosci 22, 616–626.

Mayor, R. and Theveneau, E. (2013). The neural crest. Development 140, 2247–2251.

Naba, A. (2024). Mechanisms of assembly and remodelling of the extracellular matrix. Nat Rev Mol Cell Biol.

Naba, A., Clauser, K. R., Lamar, J. M., Carr, S. A. and Hynes, R. O. (2014). Extracellular matrix signatures of human mammary carcinoma identify novel metastasis promoters. eLife 3, e01308.

Pally, D. and Naba, A. (2024). Extracellular matrix dynamics: A key regulator of cell migration across length-scales and systems. Curr Opin Cell Biol 86, 102309.

Pang, X., He, X., Qiu, Z., Zhang, H., Xie, R., Liu, Z., Gu, Y., Zhao, N., Xiang, Q. and Cui, Y. (2023). Targeting integrin pathways: mechanisms and advances in therapy. Signal Transduction and Targeted Therapy 8, 1.

Pfaff, M., Tangemann, K., Müller, B., Gurrath, M., Müller, G., Kessler, H., Timpl, R. and Engel, J. (1994). Selective recognition of cyclic RGD peptides of NMR defined conformation by alpha IIb beta 3, alpha V beta 3, and alpha 5 beta 1 integrins. Journal of Biological Chemistry 269, 20233–20238.

Pickup, M. W., Mouw, J. K. and Weaver, V. M. (2014). The extracellular matrix modulates the hallmarks of cancer. EMBO reports 15, 1243–1253.

Pietri, T., Eder, O., Breau, M. A., Topilko, P., Blanche, M., Brakebusch, C., Fässler, R., Thiery, J.-P. and Dufour, S. (2004). Conditional β1-integrin gene deletion in neural crest cells causes severe developmental alterations of the peripheral nervous system. Development 131, 3871–3883.

Pytela, R., Pierschbacher, M. D. and Ruoslahti, E. (1985). Identification and isolation of a 140 kd cell surface glycoprotein with properties expected of a fibronectin receptor. Cell 40, 191–198.

Raab-Westphal, S., Marshall, J. F. and Goodman, S. L. (2017). Integrins as Therapeutic Targets: Successes and Cancers. Cancers 9, 110.

Rozario, T. and DeSimone, D. W. (2010). The extracellular matrix in development and morphogenesis: A dynamic view. Developmental Biology 341, 126–140.

Ruoslahti, E. and Pierschbacher, M. D. (1987). New perspectives in cell adhesion: RGD and integrins. Science 238, 491–497.

Slack, R. J., Macdonald, S. J. F., Roper, J. A., Jenkins, R. G. and Hatley, R. J. D. (2022). Emerging therapeutic opportunities for integrin inhibitors. Nature Reviews Drug Discovery 21, 60–78.

Taichman, D. B., Cybulsky, M. I., Djaffar, I., Longenecker, B. M., Teixidó, J., Rice, G. E., Aruffo, A. and Bevilacqua, M. P. (1991). Tumor cell surface alpha 4 beta 1 integrin mediates adhesion to vascular endothelium: demonstration of an interaction with the N-terminal domains of INCAM-110/VCAM-1. Cell Regulation 2, 347–355.

Testaz, S., Delannet, M. and Duband, J. (1999). Adhesion and migration of avian neural crest cells on fibronectin require the cooperating activities of multiple integrins of the (beta)1 and (beta)3 families. J Cell Sci 112 ( Pt 24), 4715–4728.

Trainor, P. A. (2005). Specification and patterning of neural crest cells during craniofacial development. Brain Behav Evol 66, 266–280.

Tucker, G. C., Duband, J. L., Dufour, S. and Thiery, J. P. (1988). Cell-adhesion and substrate-adhesion molecules: their instructive roles in neural crest cell migration. Development 103 Suppl, 81–94.

Vallet, S. D., Davis, M. N., Barqué, A., Thahab, A. H., Ricard-Blum, S. and Naba, A. (2021). Computational and experimental characterization of the novel ECM glycoprotein SNED1 and prediction of its interactome. Biochemical Journal 478, 1413–1434.

Walma, D. A. C. and Yamada, K. M. (2020). The extracellular matrix in development. Development 147, dev175596.

Yang, J. T., Rayburn, H. and Hynes, R. O. (1993). Embryonic mesodermal defects in α5 integrin-deficient mice. Development 119, 1093–1105.

